# *Magnaporthe oryzae* populations in Sub-Saharan Africa are diverse and show signs of local adaptation

**DOI:** 10.1101/2020.11.17.377325

**Authors:** Geoffrey Onaga, Worrawit Suktrakul, Margaret Wanjiku, Ian Lorenzo Quibod, Jean-Baka Domelevo Entfellner, Joseph Bigirimana, George Habarugira, Rosemary Murori, Godfrey Asea, Abdelbagi M. Ismail, Chatchawan Jantasuriyarat, Ricardo Oliva

## Abstract

Rice blast caused by *Magnaporthe oryzae* is one of the most economically damaging diseases of rice worldwide. The disease originated in Asia but was detected for the first time in Sub-Saharan Africa (SSA) around 100 years ago. Despite its importance, the evolutionary processes involved in shaping the population structure of *M. oryzae* in SSA remain unclear. In this study, we investigate the population history of *M. oryzae* using a combined dataset of 180 genomes. Our results show that SSA populations are more diverse than earlier perceived, and harbor all genetic groups previously reported in Asia. While *M. oryzae* populations in SSA and Asia draw from the same genetic pools, both are experiencing different evolutionary trajectories resulting from unknown selection pressures or demographic processes. The distribution of rare alleles, measured as Tajima’s *D* values, show significant differences at the substructure level. Genome-wide analysis indicates potential events of population contraction strongly affecting *M. oryzae* in SSA. In addition, the distribution and haplotype diversity of effectors might suggest a process of local adaptation to SSA conditions. These findings provide additional clues about the evolutionary history of *M. oryzae* outside the center of origin and help to build customized disease management strategies.

## Introduction

Rice *(Oryza sativa)* feeds billions of people around the world and represents the only source of income for millions of smallholder farmers. While Asian countries produce most of the global supply, a growing demand in Africa is driving the expansion of rice cultivation in the sub-Saharan Africa (SSA) region (Nasrin et al., 2015). Despite this trend, the SSA rice yields remain relatively low, averaging 2.2 tonnes per hectare (t/ha) against the global average of 3.4 t/ha (Norman & Kebe, 2006). This low average yield is partly because the production of rice is affected by a range of pests and diseases that reduce crop yield. In fact, recent estimates suggest that crop losses due to biotic constraints in the SSA rice sector can reach up to 30% (Savary et al., 2000). One of the most challenging diseases in the region is rice blast, caused by the ascomycetous fungus *Magnaporthe oryzae* (Gurr et al., 2011). The disease was first reported as “rice fever” in China in 1637 (Agrios, 2005), but the first identification of symptoms matching its description in the SSA region was reported in Uganda in 1922 (Small, 1922). Subsequent studies reported similar problems in Ghana in 1923, Kenya in 1924, Congo in 1932, Egypt in 1935, Madagascar, Morocco, Senegal in 1952, and South Africa in 1956 (Asuyama, 1965). Since that time, the disease has spread throughout the sub-Saharan rice agro-ecosystems and has become a nuisance to most smallholder farmers in the region (Séré et al., 2013; Hubert et al., 2015; Onaga et al., 2019).

The fungus known as *M. oryzae* is actually composed of distinct genetic groups that infect multiple cereal crops (Inoue et al., 2017), but the rice-infecting isolates appear to be restricted to a single lineage (Couch et al., 2005; Choi et al., 2013). Comparative genomic studies on *M. oryzae* have provided insight into its evolutionary history in rice (Chiapello et al., 2015; Gladieux et al., 2018; Zhong et al., 2018), pointing out to a southeast Asian origin. Latorre et al. (2020) used a combined dataset to describe at least three clonal expansions from the original recombinant population occurring in the last ~100-200 years. At least two of the four global genetic groups, most likely originating from Asia, were consistently present in SSA samples. While the study in Latorre et al. (2020) involved a limited number of SSA *M. oryzae* strains, other genetic studies relying on larger collections (Chuwa et al., 2013; Mutiga et al., 2017; Odjo et al., 2018; Onaga & Asea, 2016) hinted at some amount of variation in the SSA region, which is worth exploring. Significant pathogenic variations may potentially have implications for adaptation, and consequently, management of *M. oryzae* in Africa.

Similar to other plant pathogens, *M. oryzae’* s adaptation is frequently driven by coevolution processes involving gain and loss of genes (Yoshida et al., 2016). This is particularly important for effector genes, which modulate host immunity and therefore have strong co-evolutionary signatures. *M. oryzae* effectors show a high rate of presence/absence polymorphism linked to the activity of transposable elements (Chiapello et al., 2015; Yoshida et al., 2016). This feature is likely responsible for variation in host phenotype, since some of the *M. oryzae* effectors are recognized by immunoreceptors in the rice genome, collectively called *Pi* genes (Yoshida et al., 2009; Białas et al., 2018). To date, effector studies in *M. oryzae* have also focused on a small number of isolates mainly collected from Asia. It remains unknown whether the effectors polymorphism in SSA populations shows patterns similar to the ones in Asian populations, or has been changing in the process of adaptation to the SSA region. Thus, further interrogation of *M. oryzae* populations in SSA is required to understand the connection between effector distribution and adaptation.

In this study, we combined a genomic dataset of *M. oryzae* from Asia with the genome sequence of SSA isolates collected in eleven rice-growing countries. We discovered a highly diverse pathogen population that harbors all known genetic groups predominant in Asia. Interestingly, at least two*M. oryzae* populations in SSA show different evolutionary trajectories compared to those in Asia. The patterns of presence/absence of effector genes differ from the Asian members, which might indicates a process of adaptation to the local hosts. These findings provide additional clues about the evolutionary history of *M. oryzae* outside its center of origin and help to build customized disease management strategies.

## Material and Methods

### Collection of M. oryzae isolates

All the 42 isolates used in this study were collected from 11 African countries including Burundi (4), Kenya (2), Rwanda (3), Tanzania (9), Uganda (5), Benin (4), Burkina Faso (4), Ghana (2), Mali (2), Nigeria (4), Togo (3). Additional isolates were collected in Asia (4) and Latin America (3). Isolate metadata is listed in Table S1. Infected leaves collected from rice fields were dried and kept in filter paper at 4°C before isolation. Leaves with single lesions were placed on glass rods in Petri dishes with wet filter papers and incubated at room temperature until sporulation. The sporulating lesions were examined under a stereomicroscope and a group of conidia was aseptically transferred with a transfer needle to prune agar (PA) medium (3 pieces of prunes, 1 g yeast extract, 21 g gulaman bar, 5 g alpha-lactose monohydrate, and 1 L distilled H2O) and spores were harvested in distilled water (Meng et al., 2020). Individual germinating conidia were aseptically transferred and cultured in PA. For long-term storage, each culture was overlaid with several sterilized filter paper sections and incubated at 25°C. After 10-12 days of incubation, the colonized filter paper sections were lifted from the agar surface, placed in sterile Petri dishes, allowed to dry for 3 days at room temperature, and stored at −20°C as described by Valent et al. (1986). Isolates from Ghana, Burkina Faso, Mali, and Togo were provided in filter paper format (Table S1). All the isolates are currently curated at the International Rice Research Institute (IRRI) offices at Biosciences East and Central Africa, hosted by the International Livestock Research Institute (ILRI) based in Nairobi, Kenya.

### Generation of genomic datasets

The sequence datasets used in this study were obtained from *M. oryzae* isolates collected in the field or downloaded from public INSDC databases. Fungal growth and DNA extraction were performed as described previously (Mutiga et al., 2017). DNA quality checking was carried out using a NanoDrop 1000 instrument (Thermo Fisher Scientific) and agarose gel electrophoresis. Libraries were constructed by the Beijing Genomic Institute (Shenzhen, China) using Illumina paired-end reads with an insert size of 150 bp. Sequencing was performed on Illumina HiSeq4000 with sequencing requirements set to an average coverage of 50x to 70x and a read length of 150 bp. Sequencing quality was affirmed by the fastqc algorithm and the data were trimmed by removing low-quality sequences and adapter sequences with Trimmomatic 0.36 (Bolger et al., 2014). The whole-genome sequence of *M. oryzae* 70-15 strain, reference assembly MG8 with accession number GCA_000002495 (Dean et al., 2005), was used as the reference template for mapping using BWA–mem 0.7.17 (Li & Durbin, 2009), under default parameters. Mapped reads were sorted with Samtools 1.3.1 (Li et al., 2009). Duplicate reads were removed using the *MarkDuplicates* command and all the reads in a file were assigned to a single new read-group using the *AddOrReplaceReadGroups* command with Picard 2.7 (http://broadinstitute.github.io/picard). Single Nucleotide Polymorphisms (SNPs) for each strain were called using the *HaplotypeCaller* command implemented in the Genome Analyses Toolkit 4 (GATK4.1.6.3) (McKenna et al., 2010). Subsequently, GATK’s *GenotypeGVCFs* command was applied to genotype polymorphic sequence variants for all the strains simultaneously. Hard-filtering was then performed for the raw SNP calls using the *SelectVariants* and *Variant Filtration* functions of GATK (De Summa et al., 2017). *M. oryzae* isolates with a mapping rate of less than 80% to the above-mentioned reference strain were discarded for population genetic analyses, but all the reads were used for effector mapping. In addition, the genomic datasets for 131 *M. oryzae* isolates from a global population (Gladieux, et al., 2018; Zhong et al., 2018) were downloaded from the Sequence Read Archive (SRA, http://www.ncbi.nlm.nih.gov/sra). A summary of the dataset’s sequencing yield and coverage can be found in Table S2.

### Phylogenetic and population analysis

The phylogenetic tree was built with RAxML 8.2.9 (Stamatakis, 2014). Statistical confidence for each node was set to 1000 bootstrap runs, utilizing the general time-reversible model of nucleotide substitution with the Gamma model of rate heterogeneity. The phylogenetic tree was visualized with the *ggtree* R package (Yu et al., 2017). We also performed a phylogenetic network analysis employing the neighbor net method implemented in SplitsTree 4.16.1 (Huson & Bryant, 2006). For the population structure inference, a principal component analysis (PCA) was performed on the genlight object using the *glPCA* function. The population structure was calculated from a number of clusters (K) ranging from 2 to 8, using the discriminant analysis of principal components (DAPC) implemented in the *adegenet* R package (Jombart et al., 2010). The membership probability of each isolate and the most fitting number of clusters by Bayesian Information Criterion were also performed in *adegenet*. We used another approach to infer the optimum number of clusters by calculating the, Silhouette score in the R package factoextra (Kassambara et al., 2017). To estimate genetic variation, the proportion of genetic variance due to population differentiation and a significant departure from neutrality, we calculated the genomewide nucleotide diversity (*Pi*), Wright’s fixation index *(Fst),* and Tajima’s *D* with the variants call format (VCF) file with VCFtools V.0.1.15 (Danecek et al., 2011) by a sliding window size of 50 kb. All these analysis used the same SNP positions.

### Effector mapping, distribution, and diversity

To map candidate effectors in SSA *M. oryzae* genomes, we followed the methods and resources from Latorre et al. (2020) with some modifications. In summary, a total of 178 protein-coding genes (both virulent and avirulent) from *M. oryzae* isolates that infect rice, wheat, oat, millet, and wild grasses were used to create reference effector sequences. All the 180 *M. oryzae* genome reads, 131 from Latorre et al., (2020) and 49 from this study, were mapped to effector reference using bwa-mem 0.7.17 (Li & Durbin, 2009). *Samtools coverage* from samtools 1.10 was used to calculate the mean coverage of each gene in each of the 180 blast isolates, with the minimum read depth set at 3x. The total number of mapped reads of each gene was divided by the length of that gene in the reference (Li et al., 2009). The threshold set to determine the presence of an effector gene was 80% coverage. A binary presence/absence matrix was created (Table S3). For clustering purposes, we used a total of 75 informative effector genes that show presence/absence polymorphisms. The hierarchical clustering analysis was done using the *hclust* function under complete linkage, and the distance matrix was computed in the R package *ade4* (Dray & Dufour, 2007) with *dist.binary* function adopting the Jaccard index. The PCA and effector loading analysis were performed as described in Latorre et al., (2020).

Furthermore, the initially mapped effector bam files were converted into fastq files by samtools 1.10 and bcftools 1.10 (Li et al., 2009) for variant calling. The consensus fasta files were created for each effector gene using seqtk 1.3 (Li, 2012) (Additional Data S1).To compute for genetic diversities for each effector, we chose sequences which have zero presence of “N” or unknown nucleotide base, and heterozygous base position. The effectors were then aligned using Mafft 7.453.0 (Katoh & Standley, 2013) employing the G-INS-i strategy. The final alignments were manually curated before any test was performed. Effector diversity indices were obtained using the R package *pegas* (Paradis, 2010). The functions used were as follows: *hap.div* for haplotype diversity *(Hd), haplotype* for the haplotype from the set of sequences, and *nuc.div* for nucleotide diversity (*Pi*). The pairwise rate of synonymous and non-synonymous codon changes (Ka/Ks ratio) was calculated using KaKs_Calculator 2.0 (Wang et al., 2010) implementing theYn00 model (Yang et al., 2000).

## Results and Discussion

### SSA populations of M. oryzae are highly diverse and harbor all known Asian genetic groups

To characterize the genetic composition of Sub-Saharan Africa (SSA) populations of *M. oryzae,* we combined previous datasets (Gladieux et al., 2018; Zhong et al., 2018) with the genome sequences of novel isolates collected in SSA rice-growing areas. The assembled genomes represent disease outbreaks that occurred across 13 different countries between 2012 and 2018 (Table S1). We identified a total of 66,744 SNPs among the global population. Based on the analysis of 164 global genomes (using isolates with more than 80% read mapping rate to the *MG8* reference assembly), we found that SSA populations are more diverse than earlier perceived (Latorre et al., 2020), and harbor all genetic groups previously reported in Asia (Figure 1). To reconstruct the phylogenetic signal of SSA strains, we used a maximum-likelihood analysis and found diverse ancestry (Figure 1A), where SSA genomes showed a genetic distribution consistent with multiple origins. Principal component analysis (PCA) clearly identified four distinct genetic clusters (Figure 1B), which were also confirmed by the Bayesian Information Criterion (BIC) and Silhouette score analysis (Figure S1B-C). Phylogenetic networks using Splitree (Figure S2) were consistent with reports from Latorre et al. (2020) where SSA genomes fall under the three clonal lineages, consistent with genetic groups 2, 3, and 4; and an additional highly diverse cluster representing the group 1 (Latorre et al., 2020). Population structure analysis (K = 2 > 8) using 164 global *M. oryzae* genomes also supports the hypothesis that SSA populations are highly diverse (Figure 1C) and resemble most of the diversity found in Asia (Chuwa et al., 2013; Onaga & Asea, 2016; Gladieux et al., 2018; Zhong et al., 2018; Latorre et al., 2020), representing genetic groups 1, 2, 3, and 4.

**Figure 1.**
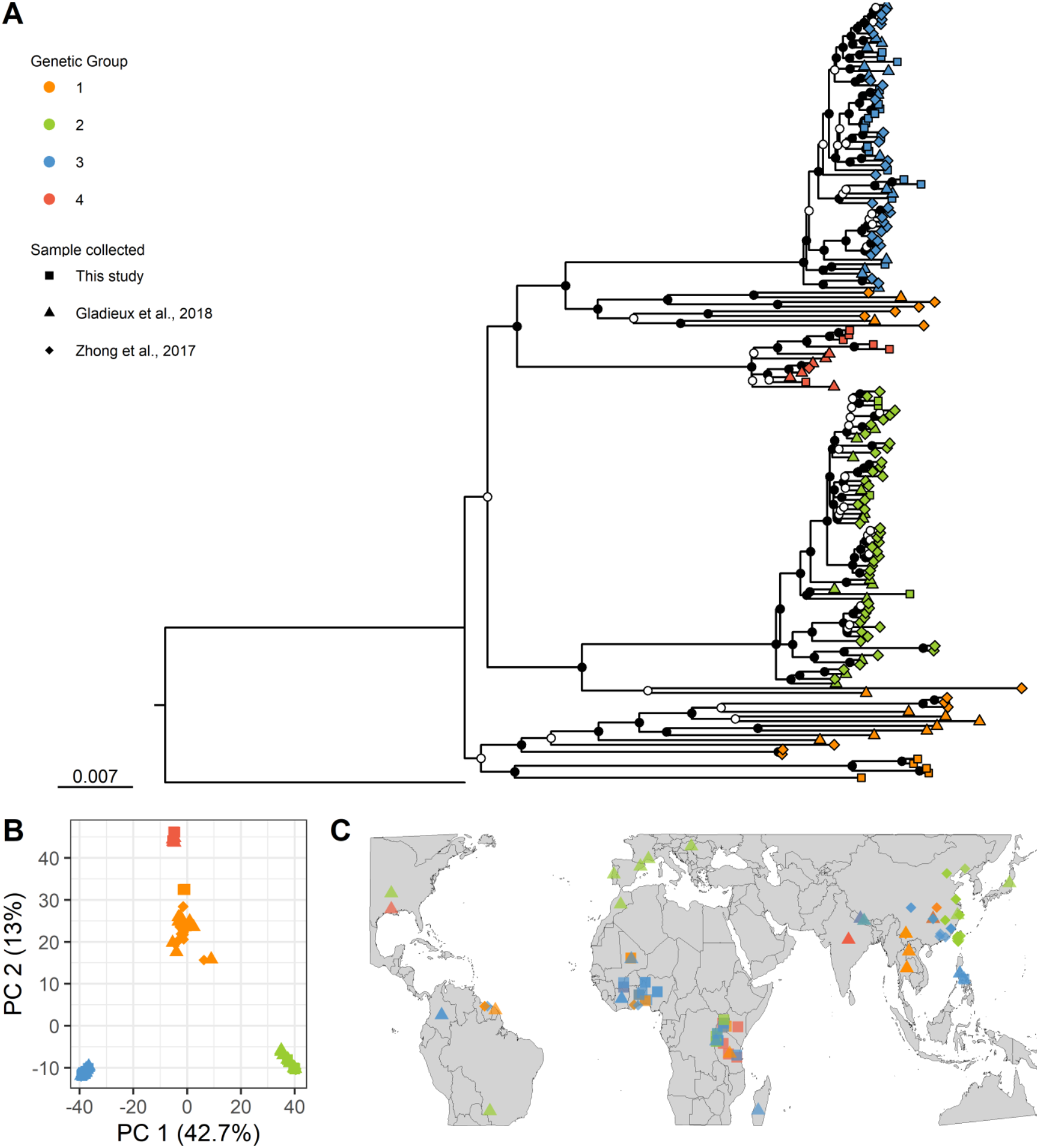
The genetic composition of rice-infecting populations of *Magnaporthe oryzae* in Sub-Saharan Africa (SSA) involves all know genetic groups from Asia. (A) A maximum-likelihood phylogenetic tree constructed with single nucleotide polymorphism alignment from 164 global isolates. Black nodes depict bootstrap scores greater than 90%. Genetic groups 1, 2, 3, and 4 were based from the results of Figure 1B and Figure S1A-C. The origin of the genomic dataset is indicated in colors. (B) A two dimensions principal correspondence analysis (PCA) showing the distribution of the genetic variability in the *M. oryzae* datasets. (C) Spatial distribution of *M. oryzae* genetic groups collected in rice-growing regions across the globe.

Recent studies suggest that the rice-infecting lineage of *M. oryzae* originated in Asia, most likely from populations infecting foxtail millet (Chiapello et al., 2015; Gladieux, Condon, et al., 2018 b). It is also suggested that this lineage emerged from a recombinant population with distinct genetic backbones (Thierry et al., 2020; Latorre et al., 2020). The rich history of rice cultivation in Africa might have allowed the early establishment of *M. oryzae* groups in different waves (Choi et al., 2019). We speculated that the first wave started before the proposed clonal expansion (Latorre et al., 2020), probably from a recombinant population similar to genetic group 1. We found representative genomes in East Africa (Uganda) and West Africa (Mali, Togo, and Nigeria), which show genetic features of the Asian metapopulation but are quite distant from any known genome in Asia. Population structure analysis of group 1 also suggests some level of sub-structuring (Figure S1A; K= 6). For instance, isolate E-UGD-32 from Tilda irrigation scheme in Uganda has unique features that might represent the emergence of novel variants in SSA. The significant differences of the SSA-1 genomes might point out to ancestral colonization rather than a recent one. A second wave appears to be more recent, involving the clonal groups 2, 3, and 4. The fact that blast symptoms were reported in Africa by 1922 suggests that such colonization occurred soon after the Asia expansion. This might explain why representatives of clonal genetic groups 2, 3, and 4 in SSA have a similar genetic background as the Asian counterparts (Figure 1A).

### SSA populations of M. oryzae are evolving in different patterns compared to Asia

To further investigate the overall demographic patterns of diversity in *M. oryzae*, we calculated population differentiation using genome-wide estimations of nucleotide diversity (*Pi*) and population substructure (*Fst*). Overall, the *Pi* values in Asia and SSA were significantly different *(p* = 0.04). While mean nucleotide diversity (*Pi*) in group 1 (*Pi* = 1.3e-04) was higher compared to the clonal groups 2 (*Pi* = 2.5e-05), 3 (*Pi* = 2.1e-05), and 4 (*Pi* = 2.45e-05) (Figure S3A; Table S4), Asia accumulate more diversity than SSA in every group (Figure 2A; Table S5). The fixation index (*Fst*) across regions (*Fst* = 0.05) or within each genetic group (*Fst* = 0.03 to 0.08) (Figure 2B) were low compared to *Fst* values between genetic groups (*Fst* = 0.25 −0.72) (Figure S4). We also detected that the major source of diversity is coming from genetic group 1 in Asia (Figure S3B; Figure S4). The observed values of nucleotide diversity and genetic structure of *M. oryzae* suggest that SSA and Asian populations belong to the same genetic pool and that each genetic group might exchange alleles between regions but not with other genetic groups. The observed patterns are consistent with panmixis in a rice-specialized asexual pathogen, but also aligns with independent drifts of each clonal lineage during the recent emergence and spread of the pathogen around the globe (Latorre et al., 2020).

**Figure 2.**
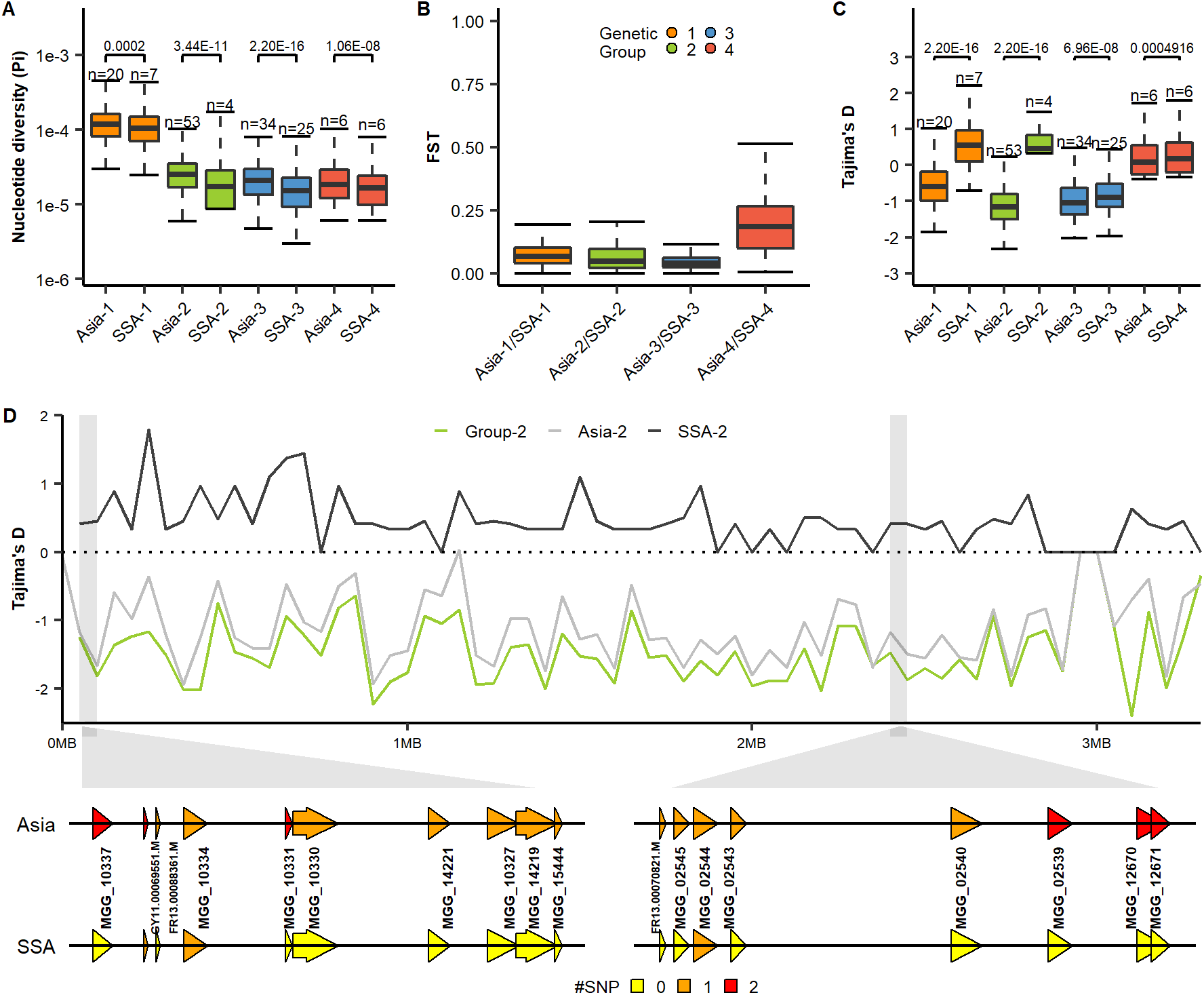
Patterns of diversity and population differentiation across rice-infecting *Magnaporthe oryzae* genetic groups in Sub-Saharan Africa (SSA) and Asia. (A) Nucleotide diversity (*Pi*) was calculated for each group. SSA1-4 and Asia1-4 represent each of the genetic groups described in Figure 1. (B) Population substructure (*Fst*) values calculated for each genetic group in SSA and Asia. (C) Genome-wide Tajima’s *D* computations of genetic groups from SSA and Asia showing substantial differences within groups 1 and 2. To compare the distribution of each genome-wide diversity test analysis in each genetic group within different regions, Mann-Whitney test was performed as shown above in the boxplot. (D) Example of the dramatic change in Tajima’s *D* values on *M. oryzae* chromosome 7 within genetic group 2 from SSA and Asia. Sequence variation of *M. oryzae* genes within a 5 kb window representing color-coded Single Nucleotide Polymorphisms (SNPs).

To explore the selection process driving the evolution of *M. oryzae* populations in Asia and SSA, we calculated genome-wide Tajima’s *D* and found significant differences at the substructure level.

Overall Tajima’s *D* values of the global *M. oryzae* showed a deviation from a neutrally evolving population (Figure S3C; Table S4), suggesting the influence of non-random events. Similar to Latorre et al. (2020), we observed negative Tajima’s *D* values (Tajima’s *D* = −0.6 to −1.2) in most of the Asian genetic groups (Figure 2C; Table S5), representing the accumulation of rare alleles in the backbone genome of *M. oryzae*. These patterns are usually indicative of population size expansion that follows a bottleneck or selective sweep. In contrast to Asia, we found SSA genetic group 1 (Tajima’s *D* = 0.56) and genetic group 2 (Tajima’s *D* = 0.58) having positive Tajima’s *D* values (Figure 2C; Table S5), which might point out to non-random removal of rare alleles from the population. Since not all the SSA groups experience significant differences in Tajima’s *D*, one hypothesis is that a sudden population contraction occurred in the SSA region, specifically affecting genetic groups 1 and 2. Interestingly, the differences in Tajima’s *D* appears to be scattered across the genome rather than concentrated in particular chromosomic regions (Figure 2D; Figure S5). For SSA genetic group 1, these differences appear to be more pronounced in chromosomes 4, 5, and 7 (Figure S5). A sudden population contraction could potentially mimic the observed Tajima’s *D* in genetic group 2, and highly likely represents the retention of prevalent populations before the contraction. However, the chromosomic Tajima’s *D* patterns observed in genetic group 1 could indicate the retention, in SSA, of cryptic or ancestral groups now disappeared from Asia.

Our data suggest that SSA and Asian populations of *M. oryzae* might be experiencing slightly different evolutionary trajectories resulting from unknown selection or demographic processes. Factors such as human interventions or weather patterns are known to produce significant selection pressure in agricultural pathogens. Recently, Thierry et al. (2020) linked climatic variation with the geographic distribution of *M. oryzae* groups. In addition, the management practices and the diversity of growing environments in Asia might be relatively different from Africa, where upland and rainfed ecologies occupy the largest share, and very little fertilizer input is used. Other factors, such as host composition, might be also likely shaping the diversity of *M. oryzae* in the region but its contribution needs further investigation.

### Effector distribution and diversification suggests local adaptation to SSA conditions

Similar to other filamentous plant pathogens, the genome architecture of *M. oryzae* appears to be shaped by the dynamics of repetitive elements (Kelkar & Ochman, 2012), which drive the rapid evolution of effector genes in response to selection (Dong et al., 2015). To understand the level of adaptation of *M. oryzae* populations in SSA, we compared the number and distribution of effector repertoires across the region and found significant differences at a substructure level. We mapped 178 predicted effector references using the genome sequence of 164 *M. oryzae* isolates. The overall number of effectors ranged from 110 to 127 per isolate and varied across genetic groups (Figure S6A-B). Similar to Latorre et al., (2020) we found a different number of effector repertoires across genetic groups. This variation can be explained by clonality and divergence processes (Figure 1; Figure 2). While the number of effectors within genetic groups 2, 3, and 4 were similar in Asian and SSA isolates, significant differences were observed within genetic group 1 (Figure 3A). It is not clear whether the differences between genetic group 1 in Asia and SSA is the result of cryptic events of genome expansion in the ancestral backbones (Dong et al., 2015) or emerged as a process of local adaptation affecting specific effectors in the region.

**Figure 3.**
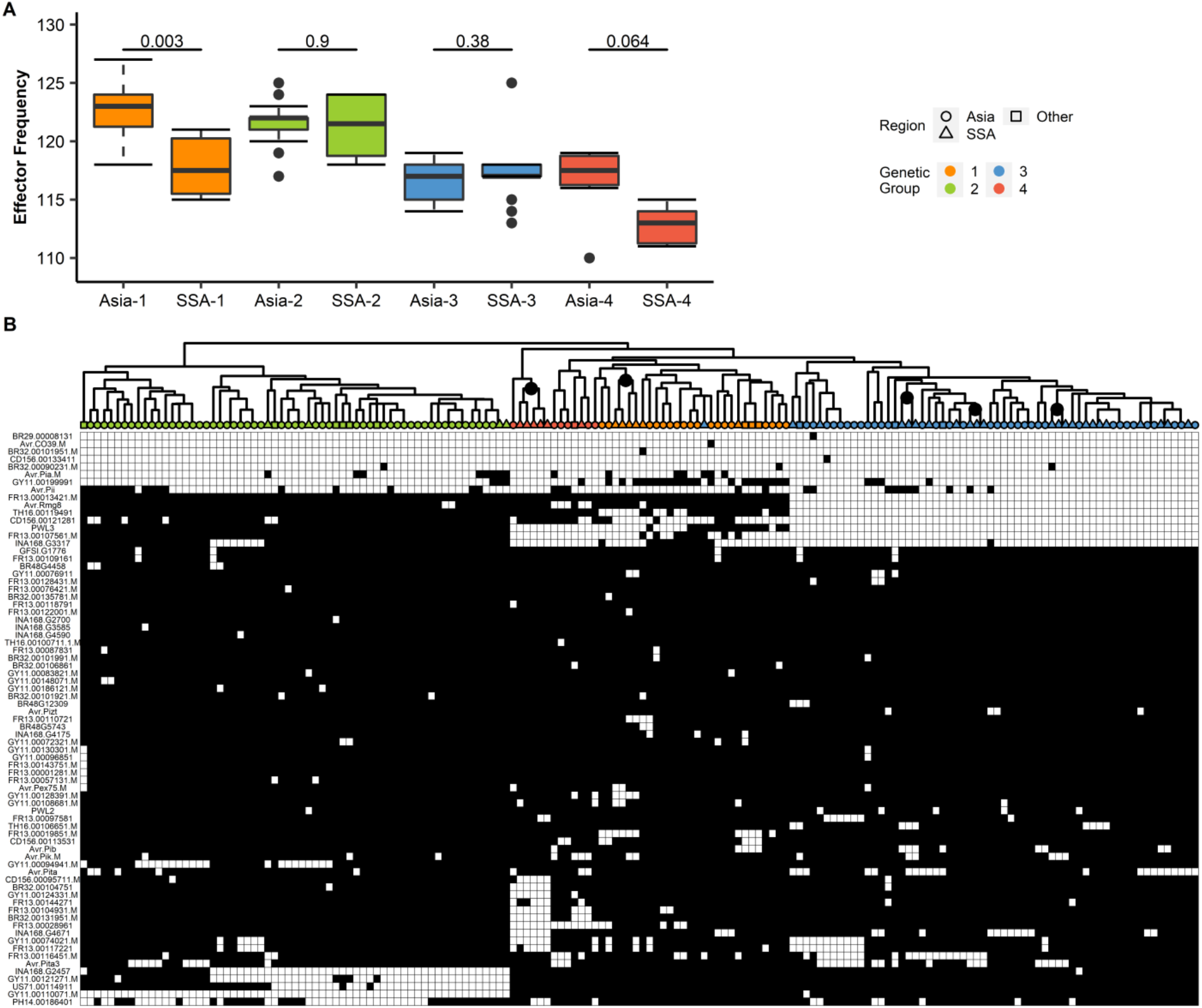
Variation in the number and distribution of candidate effector repertoires in *Magnaporthe oryzae* genetic groups collected from Sub-Saharan Africa (SSA). (A) A box plot representing the average number of effector genes in each genetic group from SSA and Asia. To compare the distribution of each effector content within the genetic group between different regions, Mann-Whitney test was performed as shown above in the boxplot. SSA1-4 and Asia1-4 represent each of the genetic groups described in Figure 1. (B) A hierarchical heatmap representing the presence/absence patterns of candidate effectors in 164 genomes. SSA (triangles), Asia (circles), and Other (squares) regions are depicted. Color labels in the tree represent each of the genetic group. Gray and white colors in the heat map represent the presence/absence of effectors as <80% of coverage. Gene names (rows) and isolate names (columns) are described in Table S1. Sub clusters in genetic groups 1, 3, and 4 are indicated as gray nodes in the tree. Complete-linkage clustering was performed as visualized in the dendrogram.

We then used a subset of 75 effectors to assess presence/absence polymorphism in the *M. oryzae* genomes (Table S3). We found that effector repertoires tend to have similar but not exact patterns in each genetic group (Figure 3B), suggesting that the clonal lineages retain virulent capabilities but rapidly gain or loss of genes from the pool. The effector loading analysis identified 16 informative effectors that explain this distribution (Figure S6C-E) and some are also present in a previous report from Latorre et al., (2020). Interestingly, we found that effector patterns from SSA *M. oryzae* genomes form specific sub-clusters within genetic groups 1, 3, and 4 (Figure 3B), which might indicate the presence of locally adapted groups in SSA. In addition, we repeated the analysis using 180 *M. oryzae* genomes and found the same distribution (data not shown).

To assess the diversity of *M*. *oryzae* effectors across regions, we extracted consensus sequences from all genes and identified allelic variants. The number of haplotypes in a subset of 96 effectors ranged from 1 to 16. The overall haplotype diversity (*Hd*), measured as the probability of finding different alleles, was higher in Asia compare to SSA for all groups. The same was also true when computing for nucleotide diversity (Figure 4A-B). We then assessed the diversity of each effector in each genetic group and found that genetic group 1 is statistically more diverse than any other genetic groups (Figure S7A-B). This observation aligned with the genome-wide diversity in Figure S3A. While more effector haplotypes were observed in Asian populations, unique effector haplotypes were present in SSA genomes (Figure S8). The presence/absence distribution and sequence differences in SSA genomes suggesting a certain level of adaptation.

**Figure 4.**
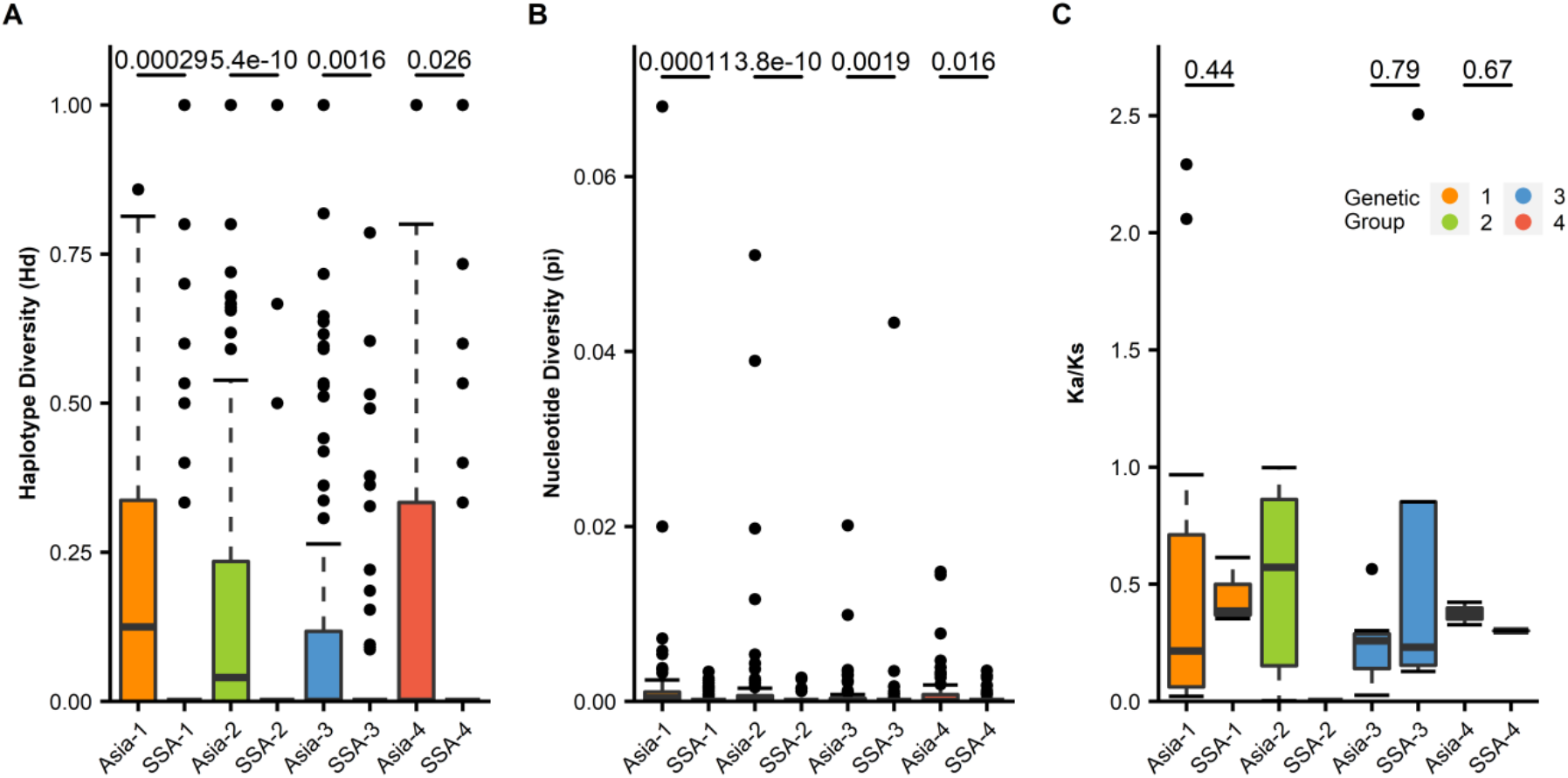
Sequence variation and signatures of selection among candidate effectors repertories in *Magnaporthe oryzae* genetic groups collected from Sub-Saharan Africa (SSA) and Asia. (A) Haplotype diversity *(Hd)* of effector genes in each genetic group from SSA and Asia. SSA1-4 and Asia1-4 represent each of the genetic groups described in Figure 1. The box plot represents only effectors with more than one haplotype. (B) Box plot representing nucleotide diversity (*Pi*) of the same dataset. (C) Distribution of synonymous and non-synonymous codon changes (average Ka/Ks ratio) across candidate effectors from *M. oryzae* genetic groups collected in SSA and Asia. To compare the distribution of each effector diversity within the genetic group between different regions, Mann-Whitney test was performed as shown above in the boxplot.

Interestingly, we only found a handful of effectors showing signatures of positive selection (Ka/Ks > 1), with not major differences between Asia and SSA (Figure 4; Figure S7C). For instance, effector PH14.00186401 showed Ka/Ks > 1 but also unique distribution of haplotypes in SSA (Figure S8). A limited number of effectors with Ka/Ks greater than 1 were also noticed by Kim et al. (2019) when comparing *M. oryzae* from different hosts. Thus, purging of allelic variants in *M. oryzae* genomes (*sensu lato*) might be a common process for this pathogen. The Ka/Ks values can be explained by purifying selection but also associated with the continuous effect of repetitive elements on the turnover of effector genes.

Altogether, our data support different evolutionary trajectories of *M. oryzae* groups in Asia and SSA. The number, distribution, and diversification of effector genes appear to indicate that host selection might have played a critical role in this process. Rice cultivation in Africa started around 3200 BP when local farmers domesticated the rice species *O. glaberrima* (Murray, 2004). While the use of *O. glaberrima* gradually declined with the introduction of the Asian species *O. sativa* (Cubry et al., 2018), it was not until 1870 when its adoption intensified (Linares, 2002). Nowadays, *O. glaberrima* has been introgressed in multiple breeding programs and it is highly represented in a significant number of released popular varieties (e.g. IRAT, ROK, or NERICA types). These varieties captured the local adaptability features from *O. glaberrima*, but might have specific stress-related genes as well. In fact, blast screenings of diverse collections of *O. glaberrima* yielded a number of resistance phenotypes (Bidaux, 1978; Silue, 1991; Yelome et al., 2018) suggesting the presence of cryptic *R* genes. If *M. oryzae* colonized SSA in multiple waves, it is highly likely that such adaption involved, at least partially, the interaction with *O. glaberrima* genotypes. The difference in host species might, in principle, explain the unique features of *M. oryzae* SSA genomes, but more detailed studies are needed to understand the broad variety of plant receptors in this African rice species.

## Conclusion

The migration of plant pathogens to new agricultural ecosystems represent a major concern for long-term strategies of food security. Understanding the events that shaped the pathogen population structure in the new setup is likely to help to develop effective control measurements. In this report, we reconstructed the evolutionary trajectory of the rice blast pathogen *M. oryzae,* present in Sub-Saharan Africa (SSA) at least for the last 100 years. The SSA populations harbor all the genetic complexity from the Asian population but display additional features that suggest local adaptation. The distribution and diversity of effector repertoires also indicate elevated virulent potential and cryptic diversity that is worth exploring in the future. The report provides important insights into the complexity of this pathogen that threatens rice cultivation in Africa and advice on the directions of future resistance management deployment strategies.

## Supporting information

All supplemental Figures

## Supplementary Material

**Table S1.** Name, origin, and year of collection of all rice-infecting *Magnaporthe oryzae* isolates used in this study.

**Table S2.** The number of mapped reads obtained by each of the *Magnaporthe oryzae* datasets used in the study.

**Table S3.** A binary matrix representing presence/absence of effectors genes in 180 *Magnaporthe oryzae* genomes worldwide.

**Table S4.** Mean and median genome-wide nucleotide diversity (*Pi*) and Tajima’s *D* of *Magnaporthe oryzae* genetic groups globally.

**Table S5.** Mean and median genome-wide nucleotide diversity (*Pi*), and Tajima’s *D* of *Magnaporthe oryzae* genetic groups in Asia and SSA.

**Figure S1.** Four genetic groups were inferred in 164 global *Magnaporthe oryzae* isolates based on the whole genome SNPs.(A) Bar plot showing the membership probability of each genome from K=2 to K=8 populations. The clusters were built using a discriminant analysis of principal components (DAPC). (B) The Bayesian information criterion (BIC), and (C) the Silhouette score both hint at 4 genetic groups as the optimum number of clusters (elbow in the BIC curve and maximum Silhouette score).

**Figure S2.** Phylogenetic network analysis of global *Magnaporthe oryzae* populations using the neighbor net method showing the four inferred genetic groups. The color denotes the four genetic groups as inferred by the PCA in Figure 1B and confirmed by the clustering analysis in Figure S1B-C.

**Figure S3.** Genome-wide genetic analysis of *Magnaporthe oryzae* genetic groups. (A) Nucleotide diversity (*Pi*) analysis of each genetic group. The *Pi* analysis presented genetic group 1 as the most diverse genetic group. (B) Fixation index (*Fst*) among genetic groups. The *Fst* analysis reveals genetic group 1 as a major source of genetic flow between genetic groups. The color in each box plot designates the pairwise comparison between genetic groups. (C) Tajima’s *D* computation among genetic groups shows negative Tajima’s *D* values. The overall patterns are similar to previous reports (Latorre et al., 2020).

**Figure S4.** Fixation index *(Fst)* between different *Magnaporthe oryzae* genetic groups across different regions. The pattern shows genetic group 1 in Asia (Asia-1) shares diversity to almost all populations. In contrast, genetic group 1 from SSA (SSA-1) shares less diversity with all the groups. Clonal lineages in Asia or SSA also show a range of differentiation. The color in each box plot designates the pairwise comparison between genetic groups in each region.

**Figure S5.** Genome-wide Tajima’s *D* from *Magnaporthe oryzae* chromosomes reveals different patterns in genetic group 1 and 2 across regions. Tajima’s *D* of each genetic group from chromosomes one to seven was built using a 50kb sliding window. The different genetic group colors correspond to the predicted groups from Figure 1B and Figure S1B-C. Tajima’s *D* values for Asia and Sub-Saharan Africa (SSA) regions are depicted as grey and black lines.

**Figure S6.** Effector repertoires in *Magnaporthe oryzae* reveal distinct patterns of diversification in each genetic group. (A) An assortment of the total number of effectors per isolate from highest (CH0333 = 127) to lowest (IN0072 =110). (B) A box plot of the total effector content in each genetic group. To compare the distribution of the total effector for every isolate in each of the four genetic groups, Mann-Whitney test was performed as shown above in the boxplot. (C) PCA biplot using 82 subsets of effectors from the presence/absence matrix (Figure 3B). The effector loading vectors are indicated in the arrow. (D) The bar plot shows the product for each effector loading vectors. The redline reveals 90% of the cumulative sum from the data, or in this case, sixteen effectors that can explain the distribution. (E) Complete hierarchical clustering dendrogram of the sixteen effectors based on the results in Figure S6E. The distance matrix was computed using the Jaccard index.

**Figure S7.** Diversity of *Magnaporthe oryzae* effectors associates with a strong purifying selection (A) Haplotype diversity *(Hd)* analysis of each effector from the four genetic groups. (B) Nucleotide diversity (*Pi*) of each effector from the four genetic groups. (C) Ka/Ks effector distribution in each genetic group. To compare the distribution of each effector diversity test analysis in each of the four genetic groups, Mann-Whitney test was performed as shown above in the boxplot.

**Figure S8.** Haplotype diversity *(Hd)* of effector repertoires in *Magnaporthe oryzae* genetic groups from Asia and Sub Saharan Africa (SSA). The heatmap shows haplotype frequency from 96 effectors present in each genetic group across regions. Effector haplotypes range from one to sixteen. There are more effector haplotypes in Asia compared to SSA, regardless of genetic group. Low frequency (yellow) and high frequency (red) is based on the total number of isolates that have that particular haplotype in each effector. The color in the text is based on the four genetic groups inferred in Figure S2.

**Additional Data S1**. The complete effector sequences files for each of the180 strains in fasta file

## Data availability

The sequences produced in this study are stored under BioProject PRJNA670311. Genomic files in Genbank format are available at NCBI. Effector sequences from each of the 180 strains are provided in Additional Data 1.

## Acknowledgment

The authors would like to thank Mr. Inosters Nzuki for technical assistance. Blast isolates from Burkina Faso, Ghana, Mali, and Togo were kindly provided by Dr. Samuel Mutiga (Biosciences eastern and central Africa – International Livestock Research Institute) to whom we are grateful. We are grateful to the DOST–Advanced Science and Technology Institute (DOST-ASTI) for free access to high-performance computing services. We also like to thank Scientists at IRRI are partially funded by the Research Program on Rice Agri-food System (RICE). Funding support also came from The Bill and Melinda Gates Foundation through the STRASA project.

## Author contribution

GO, JBDE, CJ, AI, and RO designed the research; GO, ILQ, WS, MW, performed research experiments; and analyzed the data; GO, JB, GH, RM arrange disease collection; CJ, GA, GO, RO, JB, CJ, supervised the students; RO, GO, ILQ wrote the paper

## Reference

Agrios, G. N. (2005). Plant diseases caused by fungi. Plant Pathology, 4.

Asuyama, H. (1965). Morphology, taxonomy, host range, and life cycle of Piricularia oryzae. The Rice Blast Disease, 9–12.

Białas, A., Zess, E. K., De la Concepcion, J. C., Franceschetti, M., Pennington, H. G., Yoshida, K., Upson, J. L., Chanclud, E., Wu, C.-H., & Langner, T. (2018). Lessons in effector and NLR biology of plant-microbe systems. Molecular Plant-Microbe Interactions, 37(1), 34–45.

Bidaux, J. (1978). Screening for horizontal resistance to rice blast (Pyricularia oryzae) in Africa. Rice in Africa, 159–174.

Bolger, A. M., Lohse, M., & Usadel, B. (2014). Trimmomatic: A flexible trimmer for Illumina sequence data. Bioinformatics, 30(15), 2114–2120.

Chiapello, H., Mallet, L., Guerin, C., Aguileta, G., Amselem, J., Kroj, T., Ortega-Abboud, E., Lebrun, M.-H., Henrissat, B., & Gendrault, A. (2015). Deciphering genome content and evolutionary relationships of isolates from the fungus Magnaporthe oryzae attacking different host plants. Genome Biology and Evolution, 7(10), 2896–2912.

Choi, J., Park, S.-Y., Kim, B.-R., Roh, J.-H., Oh, I.-S., Han, S.-S., & Lee, Y.-H. (2013). Comparative analysis of pathogenicity and phylogenetic relationship in Magnaporthe grisea species complex. PloS One, 8(2), e57196.

Chuwa, C. J., Mabagala, R. B., & Reuben, M. (2013). Pathogenic Variation and molecular characterization of Pyricularia oryzae, Causal agent of rice blast disease in Tanzania. Int. J. Sci and Res, 4(11), 1131–1139.

Couch, B. C., Fudal, I., Lebrun, M.-H., Tharreau, D., Valent, B., Van Kim, P., Nottéghem, J.-L., & Kohn, L. M. (2005). Origins of host-specific populations of the blast pathogen Magnaporthe oryzae in crop domestication with subsequent expansion of pandemic clones on rice and weeds of rice. Genetics, 170(2), 613–630.

Cubry, P., Tranchant-Dubreuil, C., Thuillet, A.-C., Monat, C., Ndjiondjop, M.-N., Labadie, K., Cruaud, C., Engelen, S., Scarcelli, N., & Rhoné, B. (2018). The rise and fall of African rice cultivation revealed by analysis of 246 new genomes. Current Biology, 28(14), 2274–2282.

Danecek, P., Auton, A., Abecasis, G., Albers, C. A., Banks, E., DePristo, M. A., Handsaker, R. E., Lunter, G., Marth, G. T., & Sherry, S. T. (2011). The variant call format and VCFtools. Bioinformatics, 27(15), 2156–2158.

De Summa, S., Malerba, G., Pinto, R., Mori, A., Mijatovic, V., & Tommasi, S. (2017). GATK hard filtering: Tunable parameters to improve variant calling for next generation sequencing targeted gene panel data. BMC Bioinformatics, 18(5), 119.

Dean, R. A., Talbot, N. J., Ebbole, D. J., Farman, M. L., Mitchell, T. K., Orbach, M. J., Thon, M., Kulkarni, R., Xu, J.-R., & Pan, H. (2005). The genome sequence of the rice blast fungus Magnaporthe grisea. Nature, 434(7036) 980–986.

Dong, S., Raffaele, S., & Kamoun, S. (2015). The two-speed genomes of filamentous pathogens: Waltz with plants. Current Opinion in Genetics & Development, 35, 57–65.

Dray, S., & Dufour, A.-B. (2007). The ade4 package: Implementing the duality diagram for ecologists. Journal of Statistical Software, 22(4), 1–20.

Gladieux, P., Condon, B., Ravel, S., Soanes, D., Maciel, J. L. N., Nhani, A., Chen, L., Terauchi, R., Lebrun, M.-H., & Tharreau, D. (2018). Gene flow between divergent cereal-and grass-specific lineages of the rice blast fungus Magnaporthe oryzae. MBio, 9(1).

Gladieux, P., Ravel, S., Rieux, A., Cros-Arteil, S., Adreit, H., Milazzo, J., Thierry, M., Fournier, E., Terauchi, R., & Tharreau, D. (2018). Coexistence of multiple endemic and pandemic lineages of the rice blast pathogen. MBio, 9(2).

Gurr, S., Samalova, M., & Fisher, M. (2011). The rise and rise of emerging infectious fungi challenges food security and ecosystem health. Fungal Biology Reviews, 25(4), 181–188.

Hubert, J., Mabagala, R. B., & Mamiro, D. P. (2015). Efficacy of selected plant extracts against Pyricularia grisea, causal agent of rice blast disease.

Huson, D. H., & Bryant, D. (2006). Application of phylogenetic networks in evolutionary studies. Molecular Biology and Evolution, 23(2), 254–267.

Inoue, Y., Vy, T. T., Yoshida, K., Asano, H., Mitsuoka, C., Asuke, S., Anh, V. L., Cumagun, C. J., Chuma, I., & Terauchi, R. (2017). Evolution of the wheat blast fungus through functional losses in a host specificity determinant. Science, 357(6346), 80–83.

Jombart, T., Devillard, S., & Balloux, F. (2010). Discriminant analysis of principal components: A new method for the analysis of genetically structured populations. BMC Genetics, 11(1), 94.

Kassambara, A., & Mundt, F. (2017). Factoextra: Extract and visualize the results of multivariate data analyses. R Package Version, 7(5), 337–354.

Katoh, K., & Standley, D. M. (2013). MAFFT multiple sequence alignment software version 7: Improvements in performance and usability. Molecular Biology and Evolution, 30(4), 772–780.

Kelkar, Y. D., & Ochman, H. (2012). Causes and consequences of genome expansion in fungi. Genome Biology and Evolution, 4(1), 13–23.

Kim, K.-T., Ko, J., Song, H., Choi, G., Kim, H., Jeon, J., Cheong, K., Kang, S., & Lee, Y.-H. (2019). Evolution of the genes encoding effector candidates within multiple pathotypes of Magnaporthe oryzae. Frontiers in Microbiology, 10, 2575.

Latorre, S. M., Reyes-Avila, C. S., Malmgren, A., Win, J., Kamoun, S., & Burbano, H. A. (2020). Differential loss of effector genes in three recently expanded pandemic clonal lineages of the rice blast fungus. BMC Biology, 78(1), 1–15.

Li, H. (2012). Seqtk Toolkit for processing sequences in FASTA/Q formats. GitHub, 767, 69.

Li, H., & Durbin, R. (2009). Fast and accurate short read alignment with Burrows–Wheeler transform. Bioinformatics, 25(14), 1754–1760.

Li, H., Handsaker, B., Wysoker, A., Fennell, T., Ruan, J., Homer, N., Marth, G., Abecasis, G., & Durbin, R. (2009). The sequence alignment/map format and SAMtools. Bioinformatics, 25(16), 2078–2079.

Linares, O. F. (2002). African rice (Oryza glaberrima): History and future potential. Proceedings of the National Academy of Sciences, 99(25), 16360–16365.

McKenna, A., Hanna, M., Banks, E., Sivachenko, A., Cibulskis, K., Kernytsky, A., Garimella, K., Altshuler, D., Gabriel, S., & Daly, M. (2010). The Genome Analysis Toolkit: A MapReduce framework for analyzing next-generation DNA sequencing data. Genome Research, 20(9), 1297–1303.

Meng, X., Xiao, G., Telebanco-Yanoria, M. J., Siazon, P. M., Padilla, J., Opulencia, R., Bigirimana, J., Habarugira, G., Wu, J., & Li, M. (2020). The broad-spectrum rice blast resistance (R) gene Pita2 encodes a novel R protein unique from Pita. Rice, 13(1), 1–15.

Murray, S. S. (2004). Searching for the origins of African rice domestication. Antiquity, 78(300), 1–3.

Mutiga, S. K., Rotich, F., Ganeshan, V. D., Mwongera, D. T., Mgonja, E. M., Were, V. M., Harvey, J. W., Zhou, B., Wasilwa, L., & Feng, C. (2017). Assessment of the virulence spectrum and its association with genetic diversity in Magnaporthe oryzae populations from sub-Saharan Africa. Phytopathology, 107(7), 852–863.

Nasrin, S., Lodin, J. B., Jirström, M., Holmquist, B., Djurfeldt, A. A., & Djurfeldt, G. (2015). Drivers of rice production: Evidence from five Sub-Saharan African countries. Agriculture & Food Security, 4(1), 1–19.

Norman, J., & Kebe, B. (2006). African smallholder farmers: Rice production and sustainable livelihoods. International Rice Commission Newsletter, 55, 33–44.

Odjo, T., Kassankogno, A. I., Adreit, H., Milazzo, J., Ravel, S., Gumedzoé, Y. M. D., Ouedraogo, I., Silue, D., & Tharreau, D. (2018). Diversity and structure of African populations of Magnaporthe (Pyricularia) oryzae from rice. Phytopathology (Submitted).

Onaga, G., & Asea, G. (2016). Occurrence of rice blast (Magnaporthe oryzae) and identification of potential resistance sources in Uganda. Crop Protection, 80, 65–72.

Onaga, G., Bigirimana, J., Murori, R., Cruz, C. M. V., Oliva, R., & Séré, Y. (2019). Section 4. Importance of Plant Diseases on Rice Production in Africa. In Rice diseases: Biology and selected management practices. International Rice Research Institute. http://rice-diseases.irri.org.

Paradis, E. (2010). pegas: An R package for population genetics with an integrated-modular approach. Bioinformatics, 26(3), 419–420.

Savary, S., Willocquet, L., Elazegui, F. A., Castilla, N. P., & Teng, P. S. (2000). Rice pest constraints in tropical Asia: Quantification of yield losses due to rice pests in a range of production situations. Plant Disease, 84(3), 357–369.

Séré, Y., Fargette, D., Abo, M. E., Wydra, K., Bimerew, M., Onasanya, A., & Akator, S. K. (2013). 17 Managing the Major Diseases of Rice in Africa. Realizing Africa’s Rice Promise, 213.

Silue, D. (1991). Resistance of 99 O. glaberrima varieties to blast. Int. Rice Res. Notes, 16, 13–14.

Small, W. (1922). Annual report of the government mycologist. Uganda Dept. Agr. Ann. Rept, 27–29.

Stamatakis, A. (2014). RAxML version 8: A tool for phylogenetic analysis and post-analysis of large phylogenies. Bioinformatics, 30(9), 1312–1313.

Thierry, M., Gladieux, P., Fournier, E., Tharreau, D., & Ioos, R. (2020). A genomic approach to develop a new qPCR test enabling detection of the Pyricularia oryzae lineage causing wheat blast. Plant Disease, 104(1), 60–70.

Valent, B., Crawford, M. S., Weaver, C. G., & Chumley, F. G. (1986). Genetic studies of fertility and pathogenicity in Magnaporthe grisea(Pyricularia oryzae). Iowa State J. Res., 60(4), 569–594.

Wang, D., Zhang, Y., Zhang, Z., Zhu, J., & Yu, J. (2010). KaKs_Calculator 2.0: A toolkit incorporating gamma-series methods and sliding window strategies. Genomics, Proteomics & Bioinformatics, 8(1), 77–80.

Yang, Z., Nielsen, R., Goldman, N., & Pedersen, A.-M. K. (2000). Codon-substitution models for heterogeneous selection pressure at amino acid sites. Genetics, 155(1), 431–449.

Yelome, O. I., Audenaert, K., Landschoot, S., Dansi, A., Vanhove, W., Silue, D., Damme, P. V., & Haesaert, G. (2018). Combining high yields and blast resistance in rice (Oryza spp.): A screening under upland and lowland conditions in Benin. Sustainability, 10(7) 2500.

Yoshida, K., Saitoh, H., Fujisawa, S., Kanzaki, H., Matsumura, H., Yoshida, K., Tosa, Y., Chuma, I., Takano, Y., & Win, J. (2009). Association genetics reveals three novel avirulence genes from the rice blast fungal pathogen Magnaporthe oryzae. The Plant Cell, 21(5), 1573–1591.

Yoshida, K., Saunders, D. G., Mitsuoka, C., Natsume, S., Kosugi, S., Saitoh, H., Inoue, Y., Chuma, I., Tosa, Y., & Cano, L. M. (2016). Host specialization of the blast fungus Magnaporthe oryzae is associated with dynamic gain and loss of genes linked to transposable elements. BMC Genomics, 17(1), 370.

Yu, G., Smith, D. K., Zhu, H., Guan, Y., & Lam, T. T.-Y. (2017). ggtree: An R package for visualization and annotation of phylogenetic trees with their covariates and other associated data. Methods in Ecology and Evolution, 8(1), 28–36.

Zhong, Z., Chen, M., Lin, L., Han, Y., Bao, J., Tang, W., Lin, L., Lin, Y., Somai, R., & Lu, L. (2018). Population genomic analysis of the rice blast fungus reveals specific events associated with expansion of three main clades. The ISME Journal, 12(8), 1867–1878.

