## Supplementary material for "*Magnaporthe oryzae* populations in Sub-Saharan Africa are diverse and show signs of local adaptation": All supplemental Figures

*all contribute equally

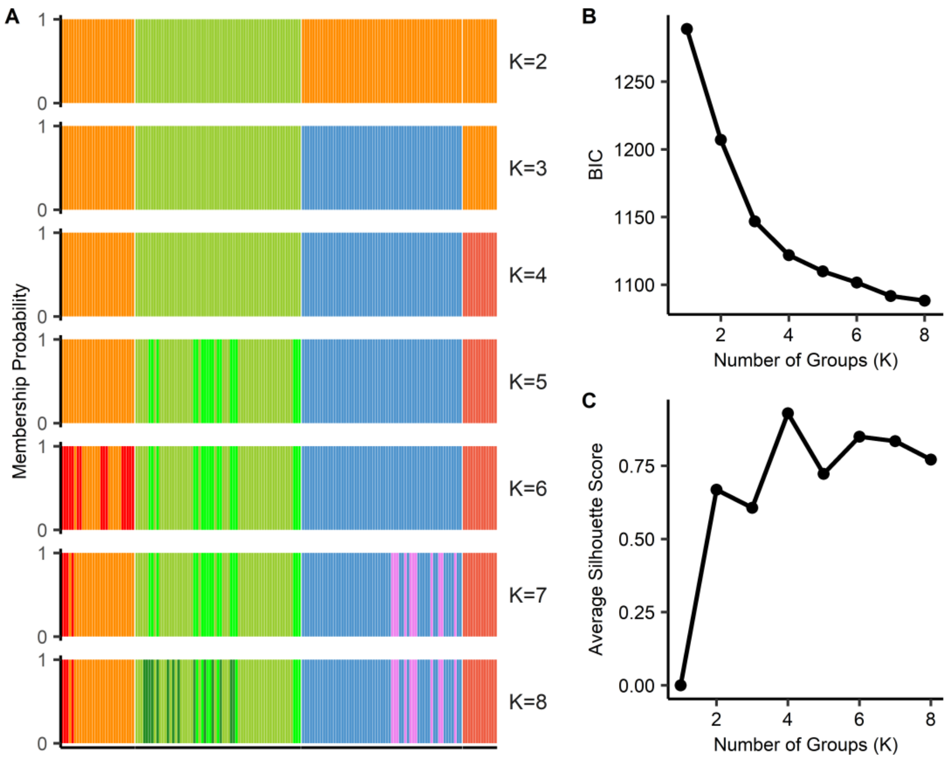


**Figure S1.** Four genetic groups were inferred in 164 global *Magnaporthe oryzae* isolates based on the whole genome SNPs. (A) Bar plot showing the membership probability of each genome from K=2 to K=8 populations. The clusters were built using a discriminant analysis of principal components (DAPC). (B) The Bayesian information criterion (BIC) and (C) the Silhouette score, both hint 4 genetic groups as the optimum number of clusters (elbow in the BIC curve and maximum Silhouette score).

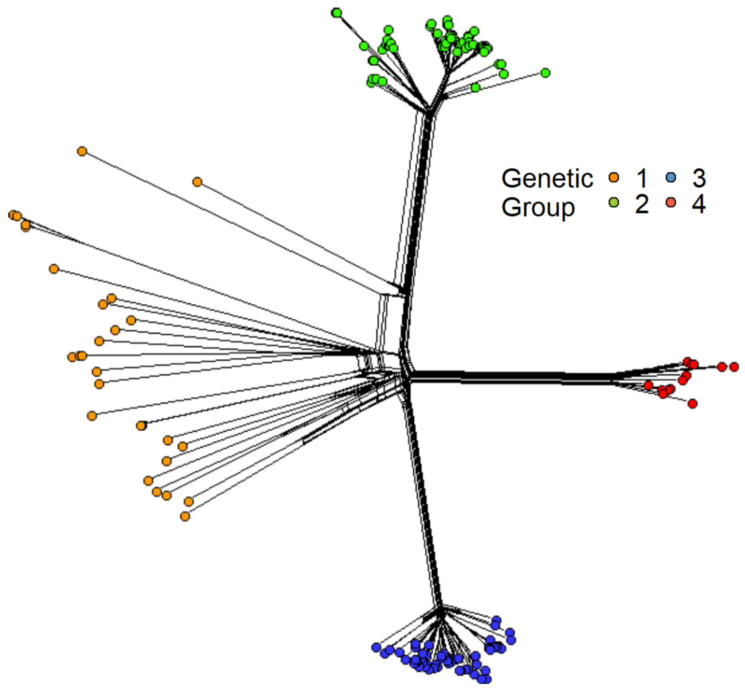


**Figure S2.** Phylogenetic network analysis of global *Magnaporthe oryzae* populations using the neighbor net method showing the four inferred genetic groups. The color denotes the four genetic groups as inferred by the PCA in Figure 1B and confirmed by the clustering analysis in Figure S1B-C.

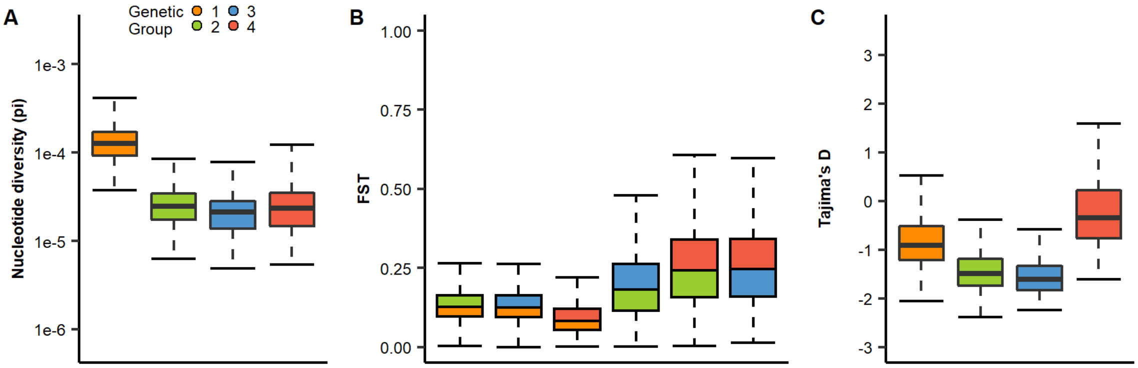


**Figure S3**. Genome-wide genetic analysis of *Magnaporthe oryzae* genetic groups. (A) Nucleotide diversity (*Pi*) analysis of each genetic group. The *Pi* analysis presented genetic group 1 as the most diverse genetic group. (B) Fixation index (*Fst*) among genetic groups. The *Fst* analysis reveals genetic group 1 as a major source of genetic flow between genetic groups. The color in each box plot designates the pairwise comparison between genetic groups. (C) Tajima's *D* computation among genetic groups shows negative Tajima's *D* values. The overall patterns are similar to previous reports (Latorre et al., 2020).

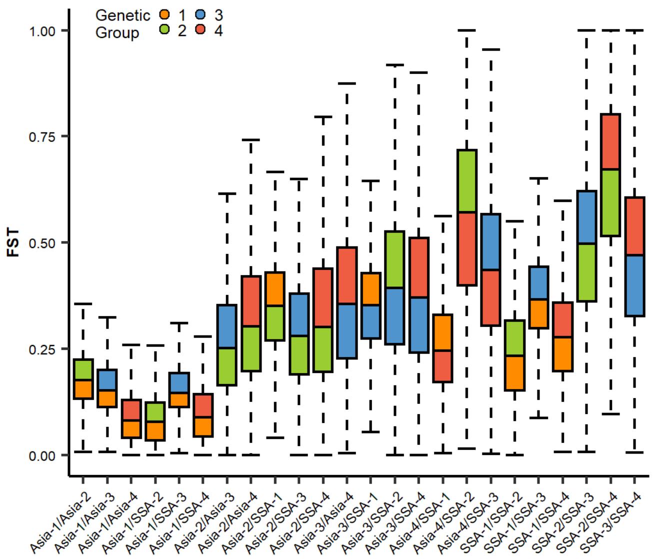


**Figure S4.** Fixation index (*Fst*) between different *Magnaporthe oryzae* genetic groups across different regions. The pattern shows genetic group 1 in Asia (Asia-1) shares diversity to almost all populations. In contrast, genetic group 1 from SSA (SSA-1) shares less diversity with all the groups. Clonal lineages in Asia or SSA also show a range of differentiation. The color in each box plot designates the pairwise comparison between genetic groups in each region.

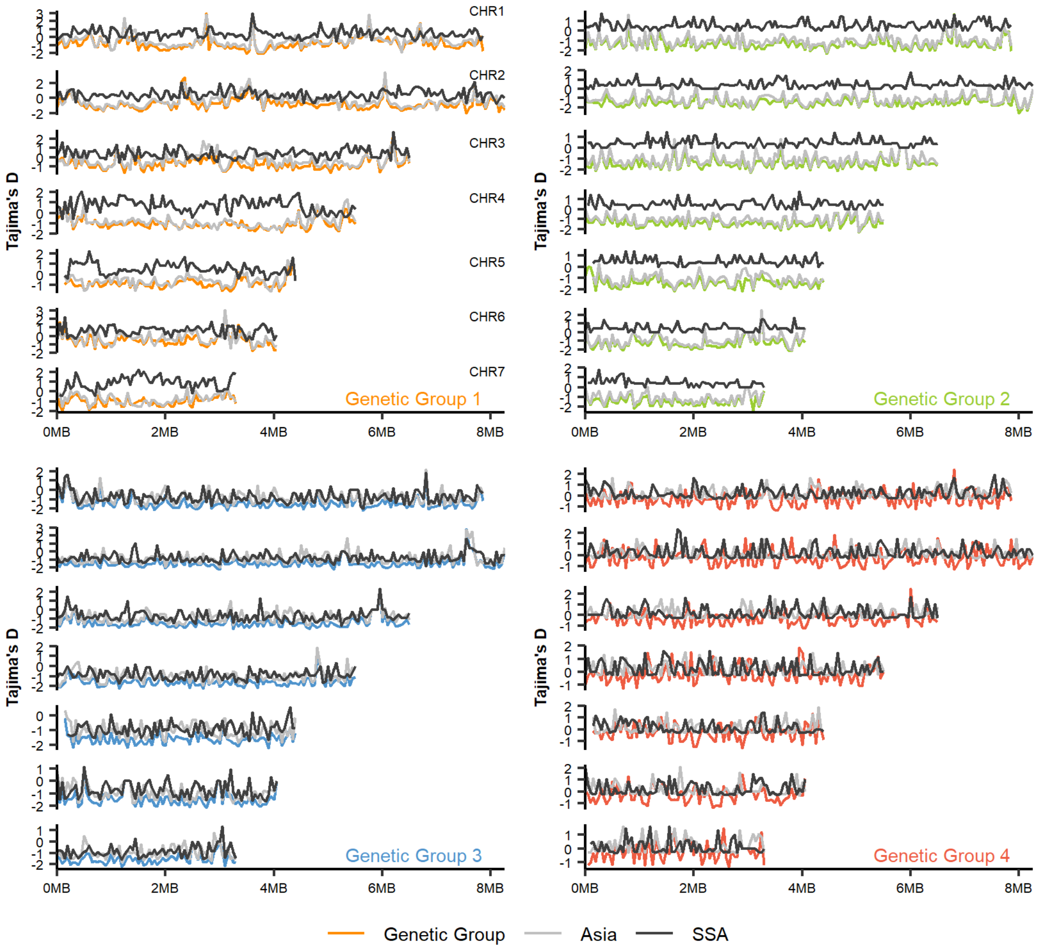


**Figure S5.** Genome-wide Tajima's *D* from *Magnaporthe oryzae* chromosomes reveals different patterns in genetic group 1 and 2 across regions. Tajima's *D* of each genetic group from chromosomes one to seven was built using a 50kb sliding window. The different genetic group colors correspond to the predicted groups from Figure 1 and Figure S2. Tajima's *D* values for Asia and Sub-Saharan Africa (SSA) regions are depicted as grey and black lines.

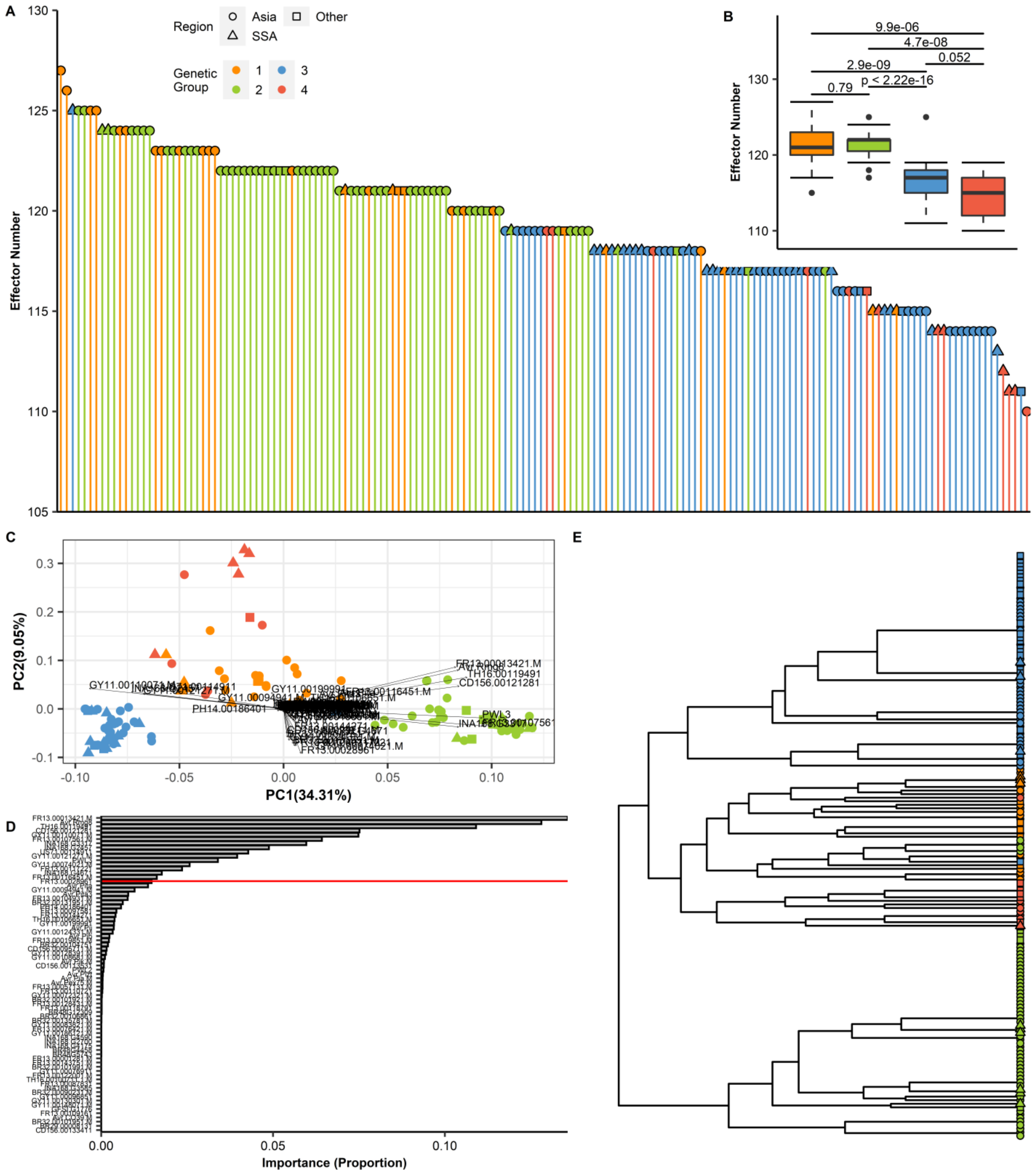
A
Effector Number
130
125
120
115
110
105
Region
Asia
Other
SSA
Genetic Group
1
2
3
4
B
Effector Number
130
120
110
9.9e-06
4.7e-08
0.052
2.9e-09
p < 2.22e-16
0.79
C
PC2(9.05%)
PC1(34.31%)
0.3
0.2
0.1
0.0
-0.1
-0.10
-0.05
0.00
0.05
0.10
FR13.00013421.M
TH16.00119491
CD156.00121281
PWL3
D
Importance (Proportion)
0.00
0.05
0.10
E

**Figure S6.** Effector repertoires in *Magnaporthe oryzae* reveal distinct patterns of diversification in each genetic group. (A) An assortment of the total number of effectors per isolate from highest (CH0333 = 127) to lowest (IN0072 =110). (B) A box plot of the total effector content in each genetic group. To compare the distribution of the total effector for every isolate in each of the four genetic groups, Mann-Whitney test was performed as shown above in the boxplot. (C) PCA biplot using 82 subsets of effectors from the presence/absence matrix (Figure 3B). The effector loading vectors are indicated in the arrow. (D) The bar plot shows the product for each effector loading vectors. The redline reveals 90% of the cumulative sum from the data, or in this case, sixteen effectors that can explain the distribution. (E) Complete hierarchical clustering dendrogram of the sixteen effectors based on the results in Figure S6E. The distance matrix was computed using the Jaccard index.

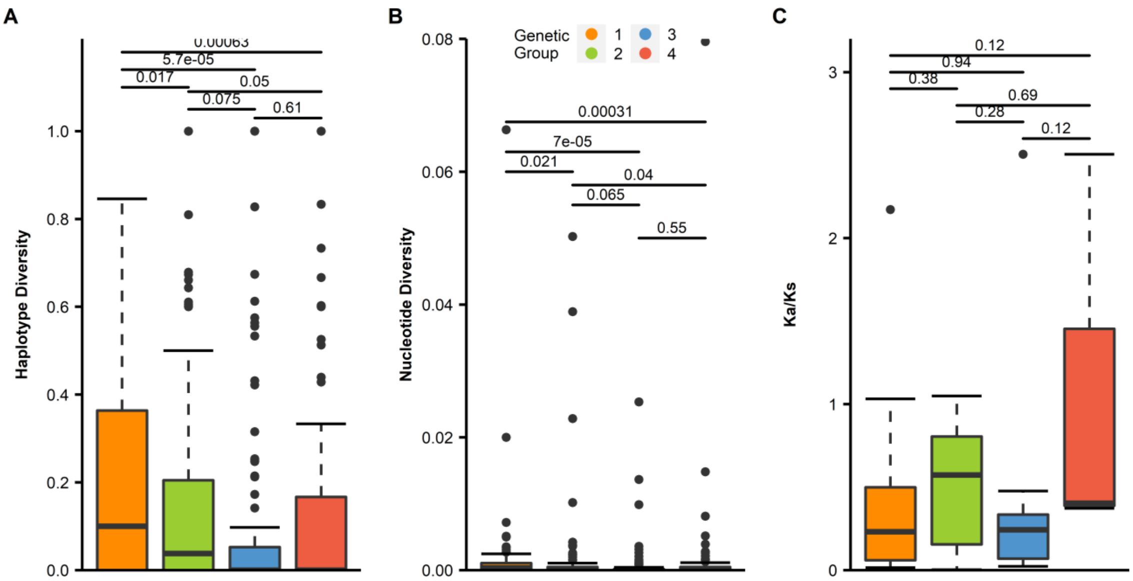


**Figure S7.** Diversity of *Magnaporthe oryzae* effectors associates with a strong purifying selection (A) Haplotype diversity (*Hd*) analysis of each effector from the four genetic groups. (B) Nucleotide diversity (*Pi*) of each effector from the four genetic groups. (C) Ka/Ks effector distribution in each genetic group. To compare the distribution of each effector diversity test analysis in each of the four genetic groups, Mann-Whitney test was performed as shown above in the boxplot.

.

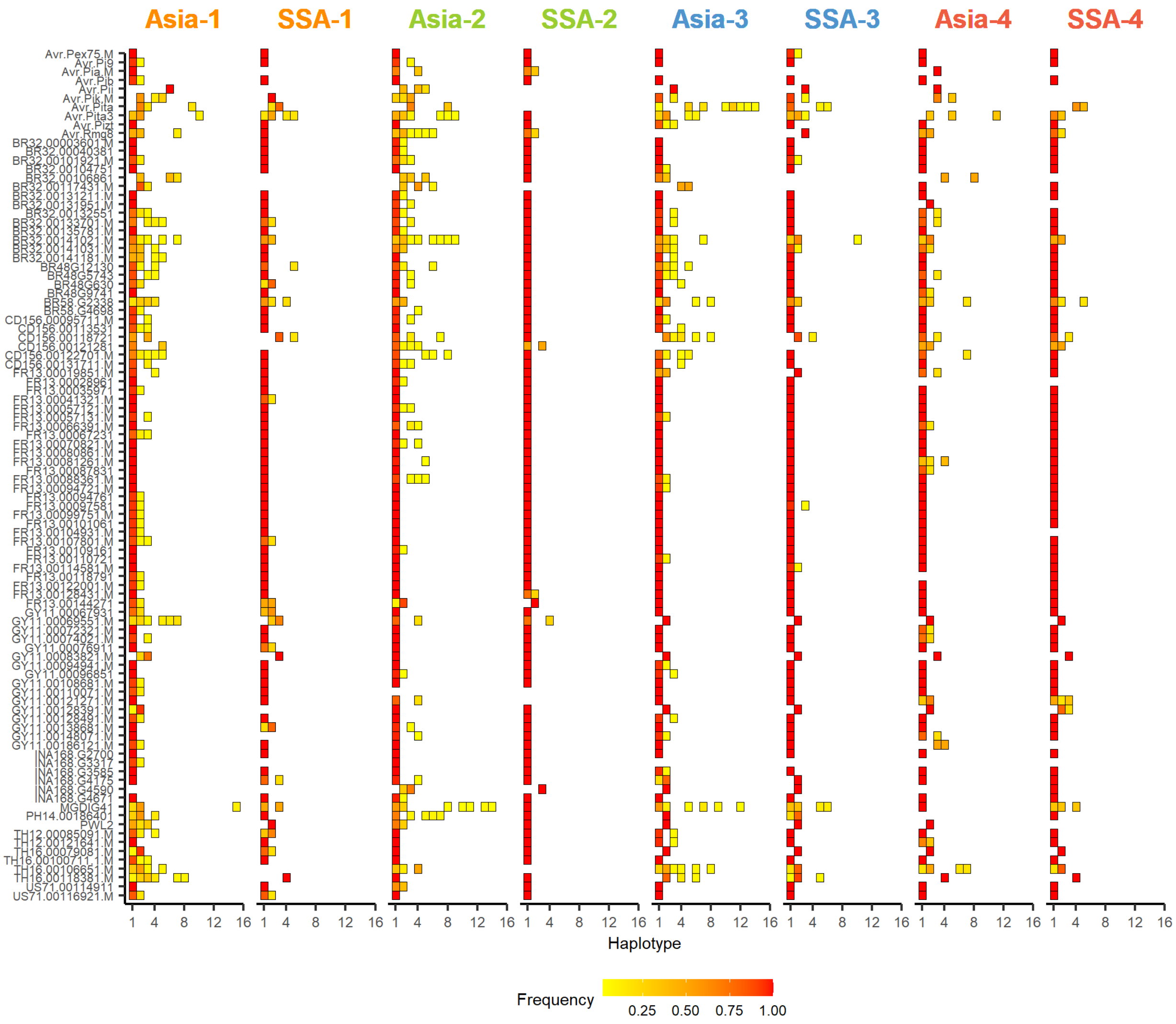


**Figure S8.** Haplotype diversity (*Hd*) of effector repertoires in *Magnaporthe oryzae* genetic groups from Asia and Sub Saharan Africa (SSA). The heatmap shows haplotype frequency from 96 effectors present in each genetic group across regions. Effector haplotypes range from one to sixteen. There are more effector haplotypes in Asia compared to SSA, regardless of genetic group. Low frequency (yellow) and high frequency (red) is based on the total number of isolates that have that particular haplotype in each effector. The color in the text is based on the four genetic groups inferred in Figure 1B and Figure S2B-C.

**Table S1.** Name, origin, and year of collection of all rice-infecting *Magnaporthe oryzae* isolates used in this study.

**Table S2.** The number of mapped reads obtained by each of the *Magnaporthe oryzae* datasets used in the study.

**Table S3**. A binary matrix representing presence/absence of effectors genes in 180 *Magnaporthe oryzae* genomes worldwide.

**Table S4.** Mean and median genome-wide nucleotide diversity (*Pi*), population divergence and Tajima's *D* of the blast genetic groups.

| | Nucleotide Diversity (*Pi*) | | *Tajima's D* | |
|---|---|---|---|---|
| Genetic Group | Mean | Median | Mean | Median |
| 1 | 0.0001262 | 0.0001208 | -0.8786072 | -0.930918 |
| 2 | 0.0000253 | 0.0000244 | -1.468771 | -1.49563 |
| 3 | 0.0000208 | 0.0000209 | -1.584732 | -1.63133 |
| 4 | 0.0000245 | 0.0000231 | -0.2561506 | -0.350067 |

**Table S5.** Mean and median genome-wide nucleotide diversity (*Pi*), and Tajima's *D* of *Magnaporthe oryzae* genetic groups in Asia and SSA.

| Genetic Group | Region | Nucleotide Diversity (Pi) | | Tajima's *D* | |
|---|---|---|---|---|---|
| | | Mean | Median | Mean | Median |
| all | Asia | 0.0001038 | 0.0001003 | -1.306864 | -1.37948 |
| all | SSA | 0.0000980 | 0.0000938 | -0.5864362 | -0.631732 |
| 1 | Asia | 0.0001170 | 0.0001145 | -0.5805084 | -0.6230595 |
| 1 | SSA | 0.0001059 | 0.0001002 | 0.5578772 | 0.510658 |
| 2 | Asia | 0.0000257 | 0.0000246 | -1.137316 | -1.178485 |
| 2 | SSA | 0.0000209 | 0.0000171 | 0.5795865 | 0.457662 |
| 3 | Asia | 0.0000213 | 0.0000204 | -0.9904772 | -1.057775 |
| 3 | SSA | 0.0000159 | 0.0000146 | -0.8631894 | -0.907495 |
| 4 | Asia | 0.0000208 | 0.0000182 | 0.1967408 | 0.0918923 |
| 4 | SSA | 0.0000171 | 0.0000158 | 0.2733068 | 0.184228 |
